# Data-Driven Design of Protein-Derived Peptide Multiplexes for Biomimetic Detection of Exhaled Breath VOC Profiles

**DOI:** 10.1101/2022.08.23.504912

**Authors:** Oliver Nakano-Baker, Hanson Fong, Shalabh Shukla, Richard Lee, Le Cai, Dennis Godin, Tatum Hennig, Siddharth Rath, Igor Novosselov, Sami Dogan, Mehmet Sarikaya, J. Devin MacKenzie

## Abstract

Exhaled human breath contains a rich mixture of volatile organic compounds (VOCs) whose concentration can vary in response to disease or other stressors. Using simulated odorant-binding proteins (OBPs) and machine learning methods, we designed a multiplex of short VOC- and carbon-binding peptide probes that detect the characteristic “VOC fingerprint”. Specifically, we target VOCs associated with COVID-19 in a compact, molecular sensor array that directly transduces vapor composition into multi-channel electrical signals. Rapidly synthesizable, chimeric VOC- and solid-binding peptides were derived from selected OBPs using multi-sequence alignment with protein database structures. Selective peptide binding to targeted VOCs and sensor surfaces was validated using surface plasmon resonance spectroscopy and quartz crystal microbalance. VOC sensing was demonstrated by peptide-sensitized, exposed-channel carbon nanotube transistors. The data-to-device pipeline enables the development of novel devices for non-invasive monitoring, diagnostics of diseases, and environmental exposures assessment.

## Introduction

In animals, the olfactory system detects and discriminates volatile organic compounds (VOCs) by binding to olfactory receptor proteins that trigger signals to the brain^1^. The smell-sensing mechanism can rapidly discriminate tiny variations in thousands of molecules at once, enabling animals to characterize their chemical environment, detect dangers, find mates, and assess food sources or toxins.

Human breath contains a rich mixture of VOCs^2^ presenting distinct VOC fingerprints that can be affected by diseases.^3^ The variations in exhaled VOC profiles can be leveraged to detect and diagnose diseases.^4,5^ Some animals can recognize these chemical signatures – for instance, dogs have been trained to detect cancers^6^ and COVID-19^7^. A small set of patient studies has identified a panel of ~ 20 VOCs in exhaled breath comprising a provisional COVID-19 fingerprint^8–11^, motivating the design of a multiplexed biomimetic sensor system to detect this and other diseases.

While an olfactory system senses VOCs through the combined action of hundreds of VOC-binding olfactory proteins^12–14^, conventional VOC sensors such as breath alcohol testers have significantly lower sensitivity and selectivity across a broad spectrum of compounds. Nonetheless, these sensors are commonplace in law enforcement and consumer markets due to their portability and simplicity, providing rapid detection at the point of use. Combining bio-inspired, combinatorial VOC detection with compact electronics could lead to the development of low-cost sensors for rapid diagnosis of COVID-19 and other respiratory infections, cancer, diabetes, and more.

Olfactory systems use odorant receptors (OR) and odorant-binding proteins (OBPs) in tandem to bind and discriminate odorants. Compared to membrane-bound ORs, OBPs are mobile aqueous VOC-transporters, showing promise as functional VOC-detecting elements^15,16^. Insect OBPs are often studied for agricultural engineering^17,18^; a dataset of insect OBP binding affinities to VOCs is compiled in Shukla et al.^19^. Peptides derived from OBPs can be effective molecular binders for VOC sensors^20^, and methods to detect peptide-to-VOC binding include quartz crystal microbalance (QCM)^21^ and piezoelectric arrays^22^. Peptide-functionalized carbon nanotube field-effect transistors (CNT-FET) are a unique sensing modality with exceptional sensitivity^23–26^.

Most reported peptide-based sensors employ a single sensing element or parent protein. Multiple aqueous peptides, however, have been used to identify wine varietals^27^, and colorimetric VOC chemosensors have taken multiplexing approaches^28,29^. For disease diagnostics, a biomimetic peptide-based biosensor could utilize multi-channel combinatorial sensing, probing for multiple VOCs in human breath. A functionalized FET sensor that transduces VOC profiles directly to electrical signals could be significantly more compact and less expensive than spectrometry instruments and would not require an expert operator. The exhaled breath VOC-based sensor would be faster and less invasive than antigen and antibody tests while possessing the specificity of biomolecular binding domains.

Machine learning (ML) approaches to COVID-19 diagnostics have applied to antigen and antibody sensing^30^; symptoms and imaging^31^; and VOC analysis by gas chromatography and mass spectroscopy (GC/MS)^9^. No multiplexed VOC analysis using biological binding agents has yet been reported. And while ML can untangle the mechanisms of biological smell^32^, it has not been used to engineer sensors that replicate olfaction mechanisms.

This work describes a peptide multiplex approach designed to detect COVID-19. Sensors comprised of OBPs were optimized using data-driven simulation in a virtual ML test bed that maximized sensitivity and selectivity to a VOC fingerprint. Sets of VOC-binding probes were extracted from the OBPs using multi-sequence alignment (MSA), incorporated into multi-functional (chimeric) peptides, validated by QCM, and tethered to a gas-sensing CNT-FET. This demonstration of a full data-to-device pipeline provides key proofs of concept for a biomolecular electronic nose: feasible disease detection with a small number of VOC-sensing channels; VOC-specific sensing by modular, chimeric peptides; and practical transduction of VOC-binding events into electrical CNT-FET device signals. This method can be adapted for a broader range of human diagnostic and environmental monitoring uses.

## Results

### Designing an OBP Multiplex Sensor

Animal olfactory systems employ many VOC-binding proteins in parallel to identify the multiple-VOC fingerprints that comprise scents. Our approach uses a similar strategy to identify disease fingerprints in human breath by applying a machine learning (ML) classification model to the outputs of an N-plex biosensor, as illustrated in **Figure 1**. Choosing from OBPs and their binding affinities identified in Shukla et al.^19^, the first design goal was to identify *N* proteins, plus a trained model, capable of detecting the disease’s VOC fingerprint. We explored protein sets by massive random search, training potential OBP-N-plex with the goal to identify diseased individuals in a simulated population of breath VOC vectors. A final set of multiplex proteins were chosen for their aggregate performance on the task.

**Figure 1:**
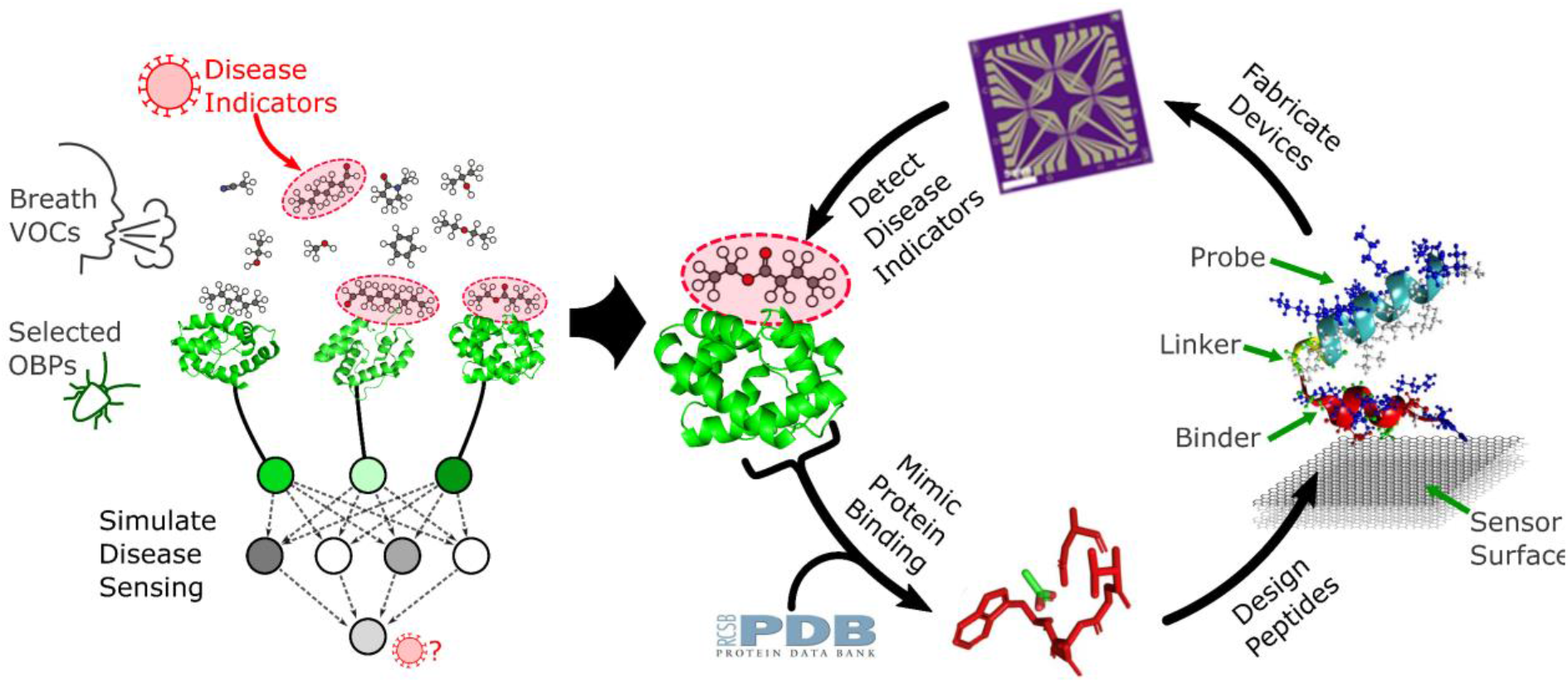
Design of a biomimetic multiplex electronic nose. OBPs were selected to allow machine learning algorithms to detect a disease’s VOC fingerprint. Peptide probe sequences that mimic OBP-VOC binding slots are integrated into modular peptides that bind to both target molecules and a molecular electronic device surface, in this case low dimension π-conjugated carbon allotropes. Finally, CNT-FET device arrays functionalized with peptides sense the VOC indicators of disease.

**Figure 2** shows simulated populations of VOC concentration vectors were generated by drawing independent and identically distributed random concentrations of VOC species found in human breath^2^, with some individuals showing elevated levels of the VOCs that comprise the disease fingerprint. A panel of *N* OBPs served as the input layer to a machine learning model, which was trained to classify samples with and without the disease fingerprint. The combined OBP multiplex and model were evaluated by its F1 score (the harmonic mean of the model’s precision and recall) on the disease detection task.

**Figure 2:**
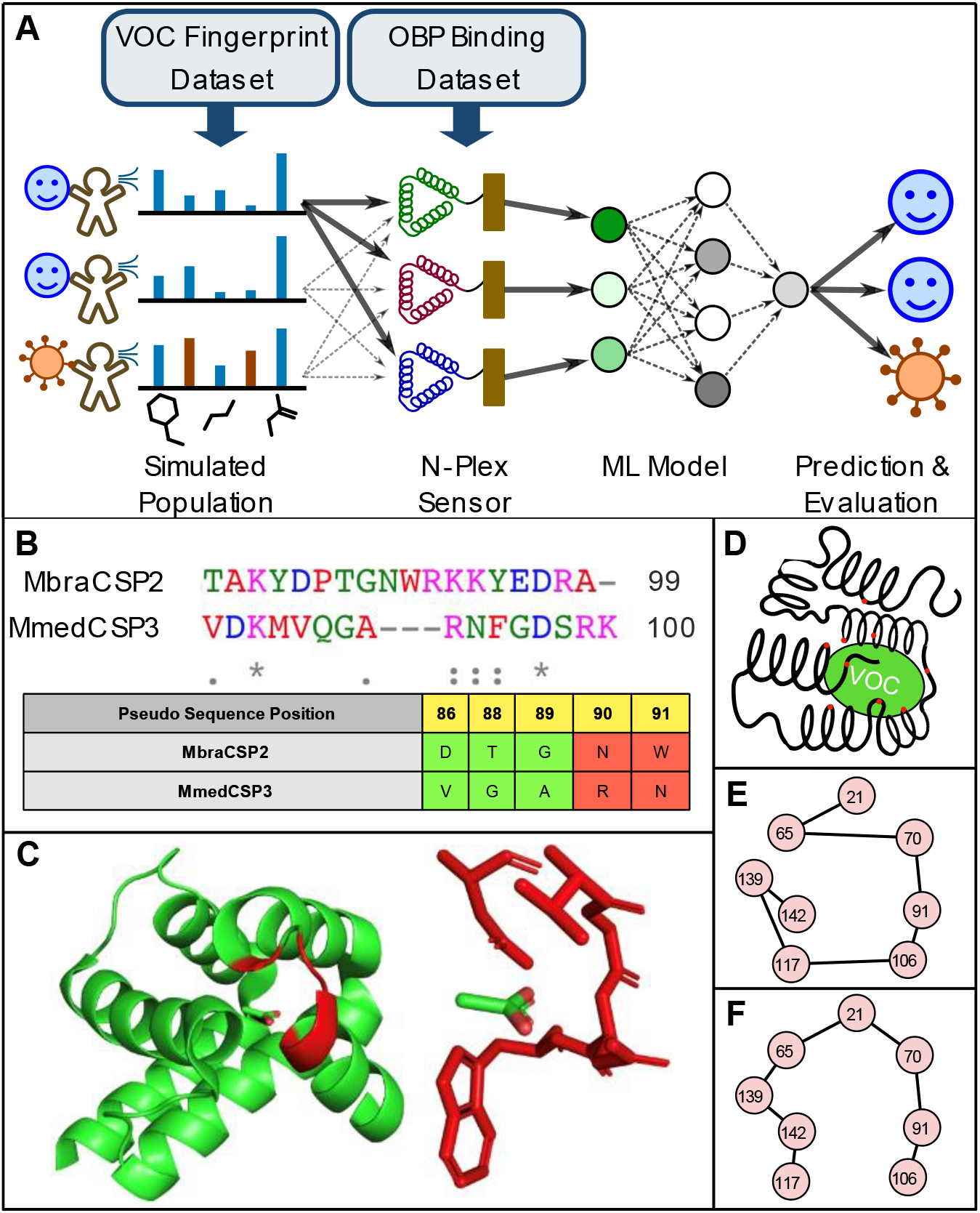
Schematic of the multiplex sensor design process. A) Simulated N-plex OBP sensors were evaluated against virtual populations of VOC profiles. Each multiplex set was evaluated on its ability to support a machine learning classification model that trained to identify healthy diseased individuals. Then, VOC probe sequences are designed from each OBP using homology matching: B) A close homological match is identified in the PDB (For OBP MmedCSP3, the matching structure is MbraCSP2, PDB ID 1KX9). C) the residues in contact with a bound ligand are identified. D) A schematic of contact residue sites spread across multiple *α*-helices. E) Sequence order is rearranged to F) traveling salesman-ordered positions to mimic residue positions in space.

Each OBP model was evaluated against the performance of a baseline “omniscient” model trained on the unaltered VOC vectors (i.e., without the lossy OBP layer). Our goal was to identify the smallest reasonable *N* where the OBP-mediated model could achieve > 90% performance relative to the omniscient baseline F1 = 0.941. In practice, *N* = 5 sensors displayed competitive performance, with the highest-performing *N* = 5 sensor achieving an F1 = 0.901 on the COVID-19 detection task, 96% of the omniscient baseline score. This OBP 5-plex was comprised of the proteins BhorOBPm2^33^, AfasOBP11^34^, MmedCSP3^35^, CpalOBP6^36^, and HcunPBP3^37^.

The search was repeated with individuals uniformly randomly assigned to be healthy, COVID-19 positive, or one of six other VOC signatures indicating respiratory diseases: rhinovirus, influenza, streptococcus pneumoniae, chronic obstructive pulmonary disease (COPD), lung cancer, and asthma. With these confounding diseases present, the omniscient model achieved an F1 = 0.687. An *N* = 10 sensor was identified that achieved F1 = 0.622, 91% of the baseline score. This sensor was comprised of the 5 COVID VOC-detecting OBPs, plus MmedOBP8^38^, LmigOBP4^39^, LstiPBP1^40^, LstiGOBP1 Val14A^41^, and AlinCSP2^42^. The 10 selected OBPs, their target VOCs, and the associated disease markers are shown in **Figure 3**.

**Figure 3.**
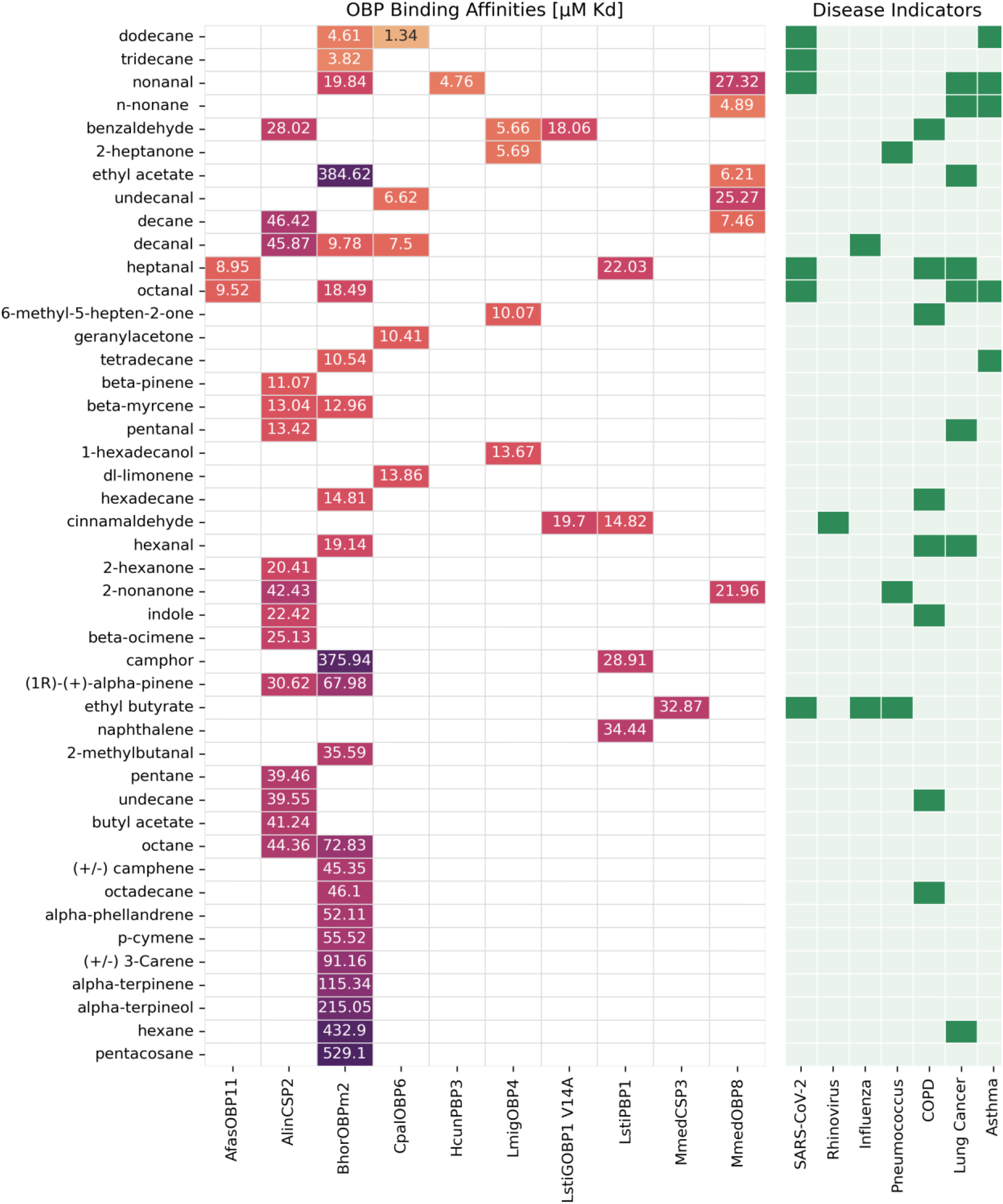
OBP-VOC binding affinities for the 10 OBPs selected to detect SARS-CoV-2 infection in the presence of six confounding respiratory disease signatures. The displayed VOCs have known binding affinity to one of the 10 OBPs and are commonly observed in human breath; the compounds are sorted with VOCs having the highest OBP binding affinity (lowest K_D_) at the top. VOCs that are disease markers but do not bind to one of the 10 OBPs are omitted.

### Probe Design by Sequence Alignment

OBPs are large molecules and are relatively difficult to synthesize, and their molecular size may limit the density of VOC binding events near the CNT and attenuate the VOC binding signal in a FET sensor, so here a short, rapidly synthesizable peptide probe sequence was extracted from each OBP that aimed to inherit the OBP’s VOC binding characteristics. By extracting the VOC-binding contact residues from each OBP as illustrated in **Figure 2**, a targeted, relatively short VOC-binding probe was integrated into a device-binding chimeric peptide.

In previous work, the peptide sequence IMVTESSDYSSY was found to self-assemble on conjugated sp^2^ surfaces, with the YSSY motif anchoring the peptide to the substrate^43^. This YSSY binding motif was used as the anchoring domain for all chimeric peptides, with a 3-glycene spacer domain linking to the probe sequences designed above. The amino acid sequence of the peptides used to functionalize the multiplex sensor were of the form *X_L_GGGYSSY* where *X_L_* is an L-length “probe” sequence that binds to target VOCs, *GGG* is a short linker, and *YSSY* is the graphite-binding motif.

For each OBP identified in a multiplex design, a binding motif was extracted from the binding slot of the protein by referencing homologically matching structures in the RCSB Protein Database (PDB)^44^. A closely matching OBP structure was identified in the PDB using BLOSUM62 alignment scores, and *L* residues that participate in ligand binding were identified in the solved structure as in Venthur et al.^45^. These binding slot residues were ordered using a traveling salesman search in 3D space to mimic the physical arrangement of the binding residues in the native protein structure. Finally, the ordered list of positions was transferred back to the hero OBP. The *L* amino acids at the specified positions became the OBP-derived probe *X_L_*.

The amino acid sequences extracted from the 10 hero sensor OBPs, listed in **Table 1**, constitute the first generation of COVID-19 VOC-targeting peptide probes.

**Table 1.**
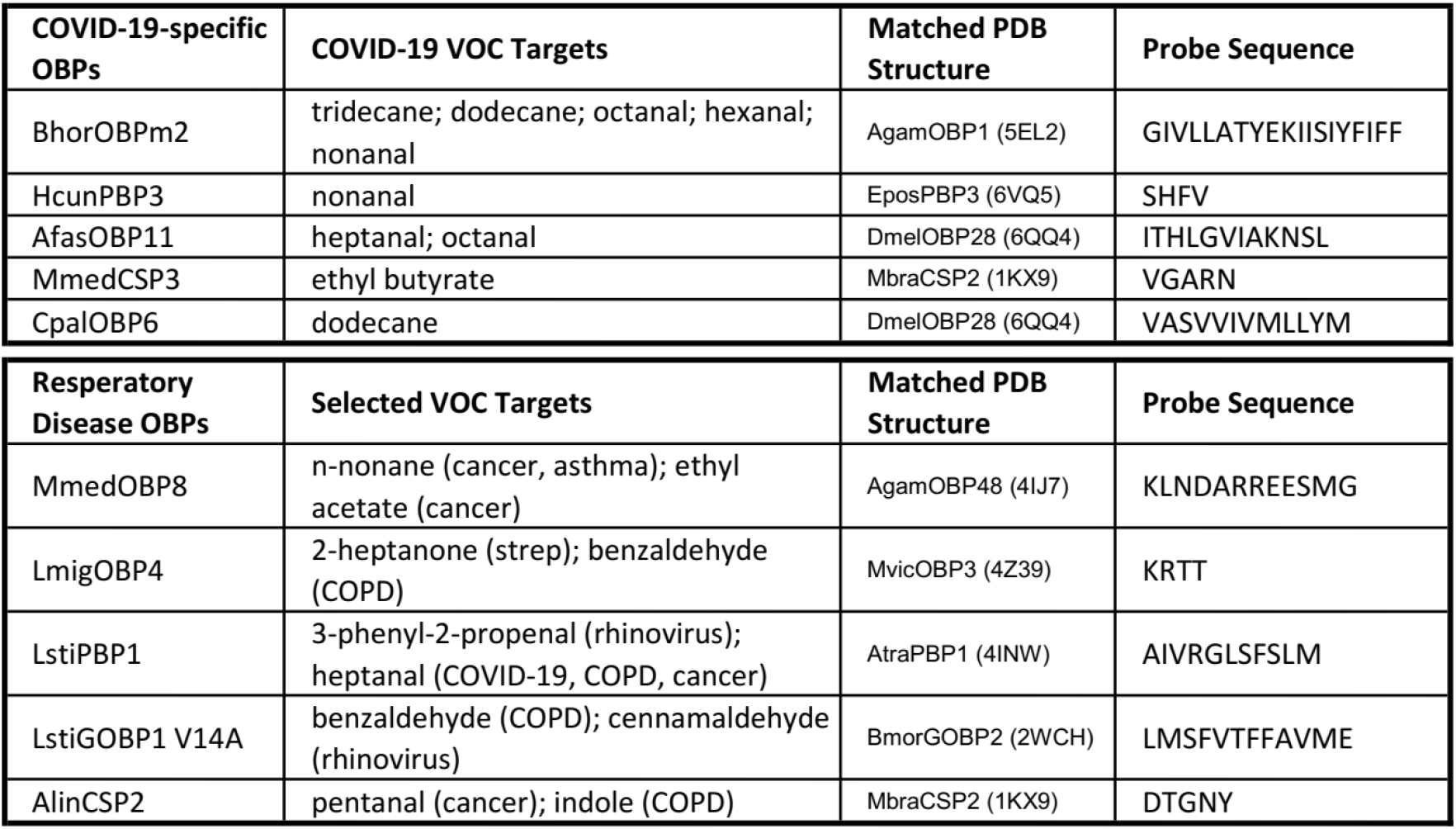
The list of OBP-derived, VOC-binding sequences in a 10-plex COVID-19-detecting sensor. Each sequence is a candidate for the “probe” portion of a chimeric peptide.

### Experimental Validation

Chimeric peptides were produced using solid-phase synthesis. To validate peptide properties, surface plasmon resonance (SPR) was used to measure the selective binding of peptides to the carbon and QCM measurements measured selective binding of probe sequences to their target VOCs versus other profile VOCs. Exposed channels of CNT-FET devices were surface treated with the two most promising peptides, and their ability to sense a target VOC was tested in a controlled gas stream. QCM and CNT-FET device evaluations were carried out in a controlled flow N_2_ environment using dry, humid, or humid VOC-spiked gas streams as depicted in **Figure 4**.

**Figure 4.**
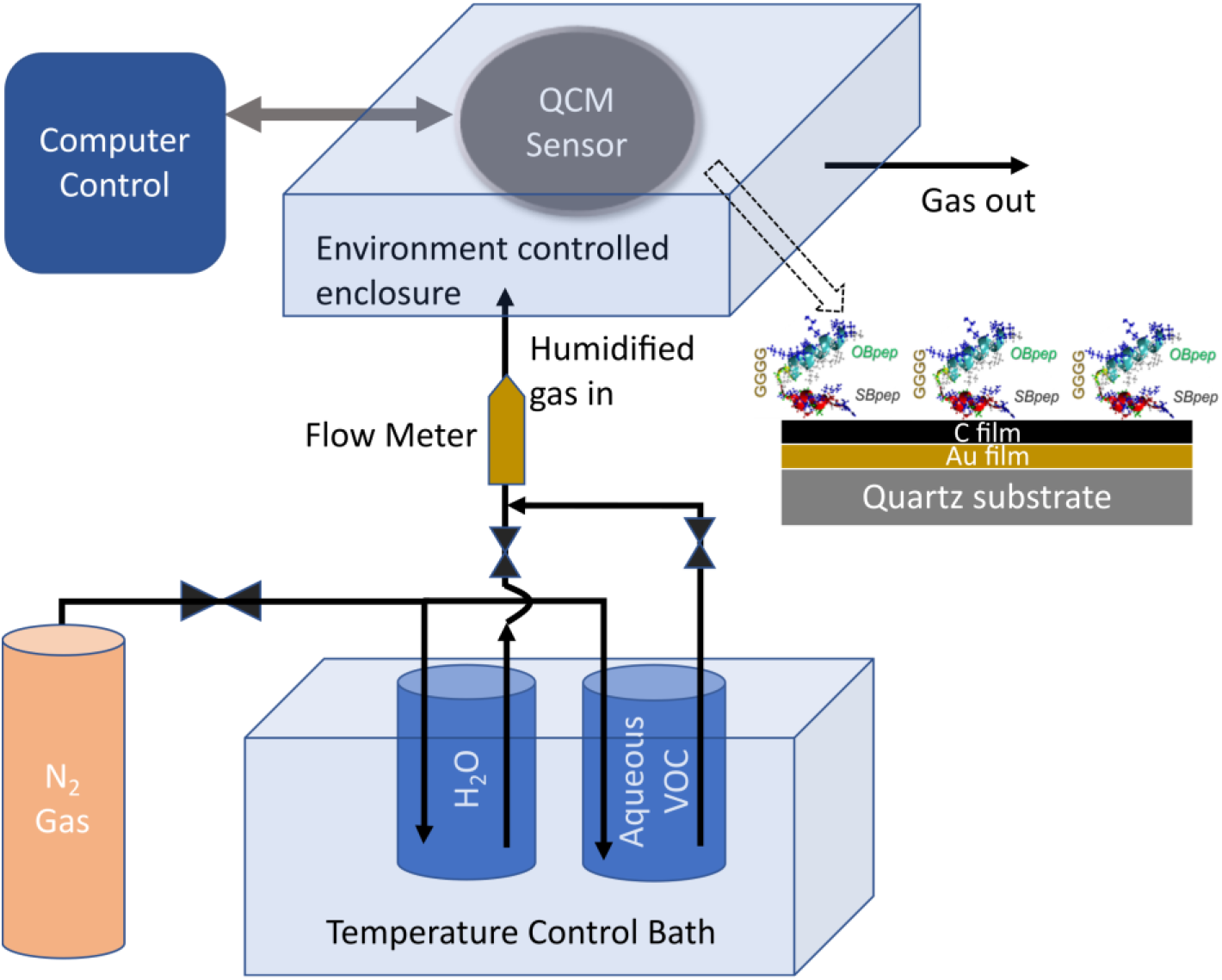
QCM experimental setup. Carrier *N*_2_ gas is humidified by bubbling it through temperature-controlled water (control) or an aqueous VOC column (test gas) and then introduced to the QCM chamber where it flows past the sensor surface.

### Chimeric Peptide Synthesis and VOC Binding Validation

Six chimeric peptides derived from the COVID-19-sensing OBPs were synthesized, see **Table 2.** SPR results revealed a greater signal response when MmedPep1-GBP was incubated on a carbon-coated gold surface than on either the bare gold or on poly-L-lysine. Signals of 158.4 ±44.1 RU were measured at 500 nM peptide concentration versus 18.4 ±1.7 RU and 6.5 ±2.9 RU for the gold and poly-L-lysine, respectively, as shown in **Figure 5**. This verified that the carbon-binding sequence YSSY of MmedPep1-GBP showed a specific preference for the carbon substrate.

**Figure 5.**
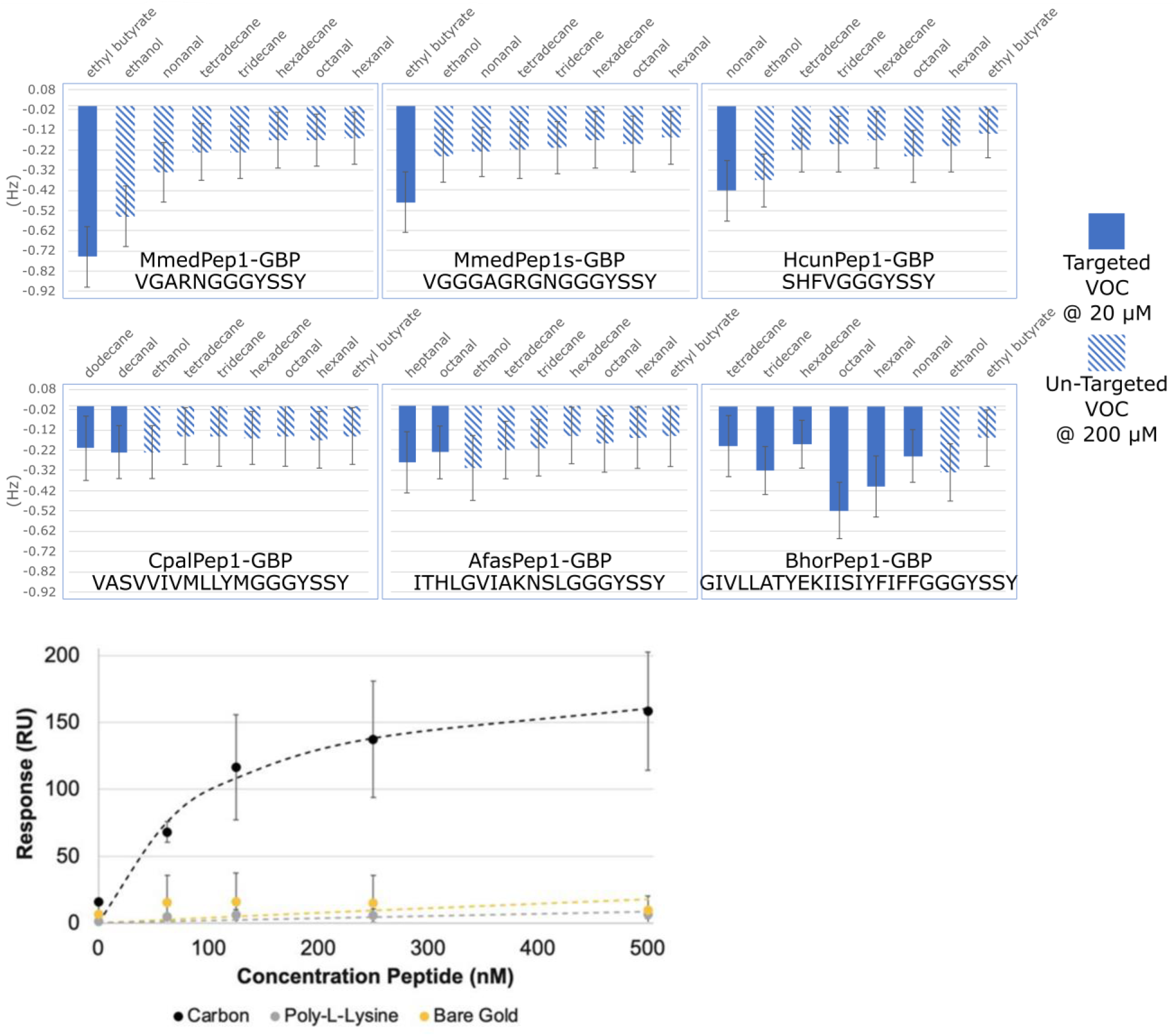
(top) Functional validation of the chimeric peptides binding to target VOCs using QCM. The flow rate of the target VOCs for each peptide is 10x lower than the confounding controls - a similar or greater reduction in QCM resonant frequency indicates binding to the target. (left) SPR data showing specific absorption of OBP3 peptide onto C-functionalized surfaces.

**Table 2.**
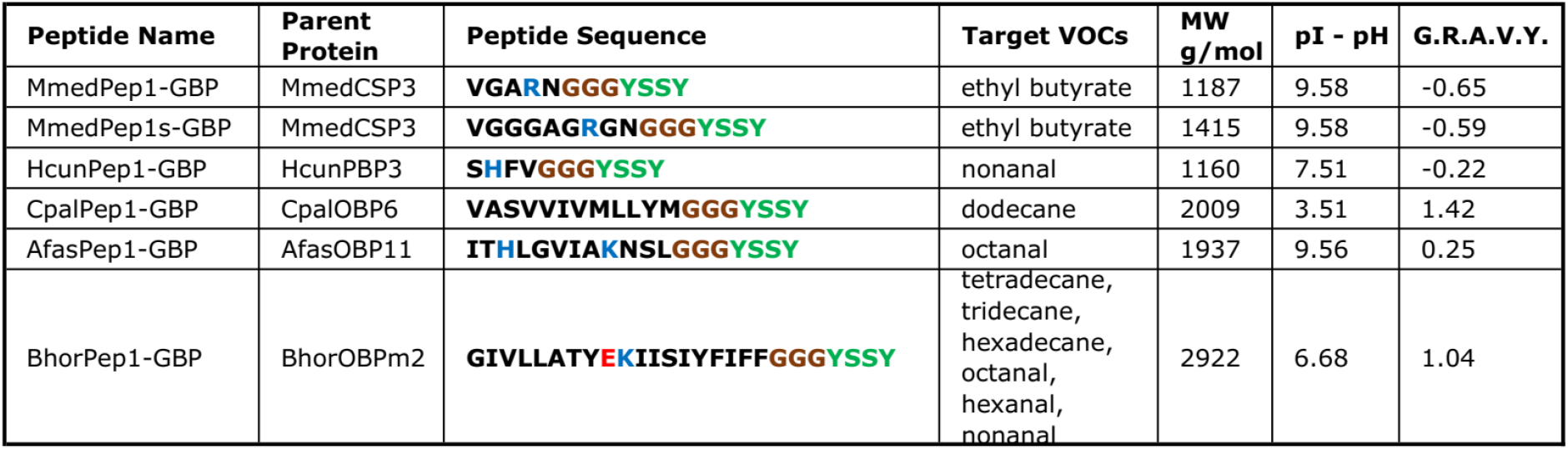
List of chimeric peptides synthesized for binding affinity validation and their critical properties. Acidic residues in the amino acid sequence are colored red and basic residues blue. Linker residues are colored brown, and the graphite-binding motif is green.

Each of the six synthesized peptides was then tested against eight VOCs. The VOCs were chosen to include the critical targeted VOCs for all synthesized chimeric peptides, plus ethanol as a common non-target. **Figure 5** shows the QCM frequency change from wet N_2_ gas to VOC-bearing wet N_2_ gas. VOC adsorption caused damping of the QCM sensor - therefore, a higher affinity surface would produce a greater reduction in resonant frequency upon exposure to the gas stream. The concentration of the target VOC for each peptide was 20 μM, 10x lower than non-targets at 200 μM. All peptides tested exhibited greater binding to their targets even at 10x lower concentration, excluding ethanol, indicating some differentiation between targeted and non-targeted VOCs. However, HcunPep1-GBP’s nonanal affinity and BhorPep1-GBP’s octanal affinity were notable and the MmedOBP8-derived peptides showed a marked preference for ethyl butyrate binding, just like their parent OBPs.

The peptide variant MmedPep1s-GBP was intended to evaluate the incorporation of glycine residues in its “probe” block to replicate the physical distances between residues in the protein’s native binding slot. This spacer variant was also strongly selective for ethyl butyrate, with reduced affinity for ethanol.

**Figure 6.**
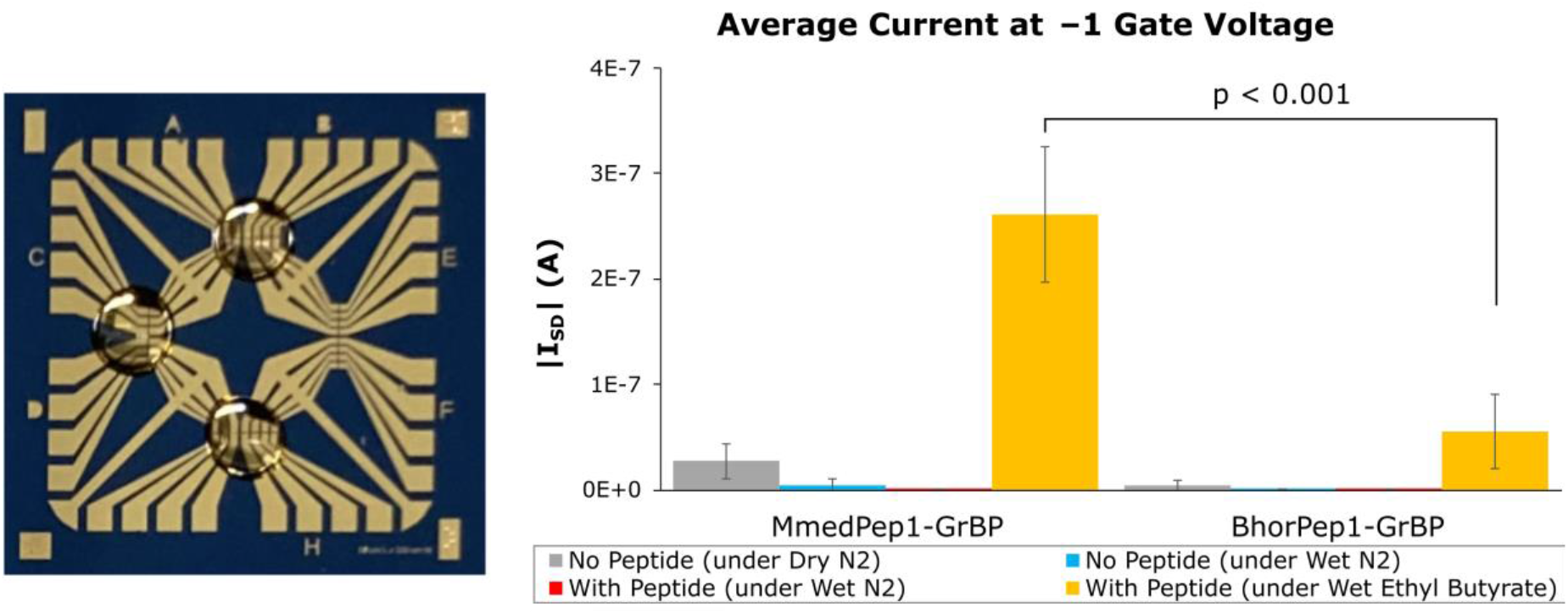
CNT-FET device fabrication and Ethyl Butyrate sensing. (a) Back gate metal oxide CNT-FET multiplexed sensor test chip during incubation of different molecular probes on three separate banks of FET sensors. (b) Steady-state source-drain currents at a gate voltage of −1V for control and peptide-functionalized CNT-FETs exposed to ethyl butyrate in flowing humidified nitrogen gas. MmedPep1-GBP was designed to be sensitive to ethyl butyrate while BhorPep1-GBP was designed to be sensitive to octanal, hexanal, nonanal and decane.

### VOC Detection by Peptide-Sensitized Carbon Nanotube Transistors

Two probes were used to incubate back gate, metal oxide CNT-FET devices to test ethyl butyrate response: MmedPep1-GBP and BhorPep1-GBP, which were predicted to bind to ethyl butyrate and a range of aldehydes and alkanes, respectively. Low source-drain currents were observed for bare CNT devices without peptide or for peptide-sensitized devices in an N_2_ gas stream when monitoring the sensing channel (200 μm length by 50 μm width) at a back gate voltage of −1 V for a 100 nm SiO_2_ gate dielectric. Flow rate was set to 45 SCCM using a flowmeter (Matheson, model FM-1050). Upon exposure to 378 mM ethyl butyrate solution (5% v/v in H_2_O), however, MmedPep1-GBP-sensitized CNT-FET measured 261.17 ± 64.18 nA of current, compared to 55.72 ± 35.14 nA for the aldehyde-sensitive BhorPep1-GBP (*p* < 0.001). This indicates that (i) the selective binding observed in QCM translated to signal transduction in a CNT-FET device, and (ii) the response was specific, contingent on the binding tendencies of the peptides on each channel.

## Discussion

Multi-channel biomimetic exhaled breath VOC sensing could be groundbreaking for public health, pandemic management, rapid disease diagnosis, and wellness monitoring. This study demonstrated a vertical design loop of such a device, from biological data to a functioning, highly specific gas sensor. With VOC and OBP datasets as inputs, we designed and validated a multiplex of VOC binding peptides for selective disease and tested on-chip the functionality of a vapor phase peptide-sensitized VOC-detecting transistor sensor. Simulated human VOC signatures were used to identify a set of OBPs capable of detecting COVID-19’s VOC signature, and probe sequences were extracted that mimic protein binding using multi-sequence alignment homology matching. Chimeric peptides were designed and synthesized with VOC- and sensor-binding domains; and SPR and QCM were used to validate the peptides’ surface- and VOC-binding credentials. Our experiments confirmed that binding residues retained their VOC-selective properties when extracted from an OBP protein and that chimeric probe peptides were readily adsorbed onto a carbon device. Finally, we functionalized back gate metal oxide CNT-FETs with the most promising peptide candidates and demonstrated electronic detection and differentiation of the VOC ethyl butyrate by a two-probe multiplex.

Combinatorial sensing of VOC fingerprints by olfactory proteins is well studied, including their poly-selectivity^12^, associated neurology^13^, and encoding structure^14^. However, these studies focus on zoological systems with hundreds of sensing proteins, adapted to detect thousands of distinct odors but impractical for transfer to a compact electronic device. When the detection target is restricted to a single disease, we demonstrated that a massive reduction of sensor complexity from hundreds of channels in the biological system to just five is possible without sacrificing device performance. Ten channels were sufficient to de-convolute multiple overlapping disease signatures within the complex matrix of VOCs in human breath using machine learning models. Because the sensor design algorithm made highly conservative assumptions, including a strictly binary sensor response and zero correlations in breath VOC concentrations besides the disease signal, we expect that a real sensor could perform to the predicted level or better.

The extraordinary parts-per-billion sensitivity of peptide-bound CNT-FETs has been explored in prior works with single-target probes based on a handful of highly characterized proteins such as ASP1^23^ or LUSH^25^, and Sim et al.^26^ selected multiple peptides based only on CNT binding affinity with a phage display library. This study is the first to demonstrate *a-priori* design of many VOC probes that together target a specific multi-VOC disease fingerprint. We have “shown all of our work” by reporting every QCM result from all synthesized peptides. All six peptides exhibited some preferential binding to their intended VOC targets, and the MmedCSP3-derived probes were highly selective for their target analyte, ethyl butyrate. Even with this small sample size, our single-shot success rate suggests that this workflow is a practical approach to sensitizing a VOC fingerprint-detecting multiplex. The probes not selected for CNT-FET testing were weakly selective for their targets, and guided evolution would likely amplify their preferential binding properties, completing the sensor - we leave these steps for a future publication.

The multiplex peptide CNT-FET sensor provides a compact and adaptable platform for development of gas phase sensors for any analyte that can be targeted for binding to a biomolecule, and we expect to see development of probe ensembles for selective diagnosis of other targets such as other respiratory infections, early-stage cancers, diabetic ketoacidosis, general health and wellness, and environmental toxins. Major enhancements to the methodology could include expansion of supporting datasets to include additional olfactory and gustatory proteins, as well as more sophisticated modeling of human VOC breath profiles. Powerful computational methods could expand the probe search space, such as deep chemoinformatic neural networks that may be able to perform direct probe-to-VOC binding affinity predictions. And atomic-scale imaging and density functional theory modeling the assembled structures formed by chimeric peptides and VOCs on CNTs could reveal the physical principles of signal transduction in peptide-CNT-FET devices. Most critically, the effect of full human or human-equivalent VOC mixtures on the precision of these devices remains to be seen. Use of high-throughput methods for peptide synthesis, VOC-binding assays, and VOC mixture testing will be critical to the development of practical, mobile sensor devices.

In conclusion, a data-driven computational method for the design of a VOC-specific OBP multiplex was able to find a compact (≤ 10 channel) sensor design that can detect disease. We demonstrated peptide molecular probes derived from OBPs that enable selective sensing of components of the COVID-19 VOC signature by sensitized carbon nanotube-based transistors. Synthesis, device binding, and signal transduction properties of the solid-binding peptide probes were validated using spiked VOC gas streams. Animal olfaction is an immensely complex system but combining biology-derived sensing elements with machine learning pattern recognition allows electronic devices to harness the molecular detection capabilities of olfactory proteins in compact, targeted sensors. These probes are computationally designed, easily synthesized, and can be rapidly integrated into compact bioelectronic devices at point-of-care. In moving past single analytes to detect the myriad VOCs that constitute scents, an abundance of public health, clinical, and environmental monitoring applications may soon be possible.

## Methods

### Simulated Populations of VOCs

One simulated breath was generated by drawing a concentration value *c* for each common breath VOC from a log-normal distribution: 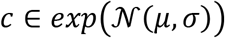. A “sick” individual is simulated by increasing the mean of the distribution of the VOCs in the COVID-19 fingerprint by a scaling factor of *v*, i.e., 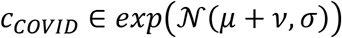. Larger *v* effectively reduces the difficulty of the diagnosis task by boosting the disease fingerprint - in practice, *v* was tuned to produce a realistic result in a baseline model. Each synthetic population comprises *M* vectors *C* = {*c*_1_, *c*_2_,... *c_V_*} of *V* VOC concentrations, with some fraction possessing elevated concentrations from the disease fingerprint. To design a multiplex sensor, the ability of a set of OBPs, in combination with a model, to classify diseased and non-diseased vectors in this population was optimized in a virtual testbed.

### Simulated OBP Sensors

Each unique OBP specifically binds to a unique subset of VOCs. To model this behavior, the testbed deployed one of two rules for OBP-VOC signal transduction. The first rule assumed nonlinear OBP transduction - that they activate in a binary manner and do not distinguish between binding events to different VOCs. An OBP with experimentally determined binding affinities 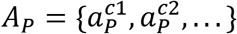 would “activate” when any *C* > *A_P_*, that is 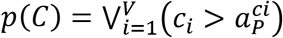. The second rule assumed that a gas mixture could be scaled linearly over time in a device and reported the “time” scaling factor at which an OBP activated - that is 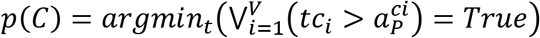. Both rules are conservative, assuming no linear response to VOC concentrations and an inability of any single OBP to differentiate between different bound VOC species. The 5-plex COVID-19 OBPs were identified using the binary VOC sensor model, and the 10-plex multi-disease sensor was identified using the time-scaling VOC model.

The first layer of each prospective disease classification model consisted of a set of *N* OBP proteins, each of which read the input VOC vector according to the binary or time-scaling activation function *p*. An *N*-plex OBP sensor measured the VOC vectors *C* from the population to produce the *N*-long sensor readings *P_OBP_* = [*p*_1_(*C*), *p*_2_(*C*), … *p_N_*(*C*)]. The N-plex would then be evaluated based on the ability of a model to classify diseased individuals using inputs *P_OBP_*.

### Machine Learning Classification

Machine learning models were implemented using adaptive boosted decision tree ensembles: SciKitLearn’s^46^ AdaBoostClassifier with DecisionTreeClassifier base estimators. Each model *f* accepts an input vector of VOC concentrations or OBP responses to VOC concentrations, generated as described above. It produces *y’,* the predicted binary disease state of the vector.

### Model Evaluation

Each multiplex sensor set *X* was evaluated on its ability to accurately train a machine learning model to classify healthy and diseased individuals in a virtual testbed. A mixed population of *M* VOC vectors is drawn 50%/50% from healthy and diseased distributions and the population is split randomly into 90%/10% train and test sets.

An “omniscient” classifier model was evaluated as a baseline. This model received an input of the true VOC concentrations *C* from the synthetic population, i.e., *f_om_*(*C*) → *y*’. Model *f_om_* trained on the training set and reported an *F*1 score, defined as *F*1 = 2(*precision · recall*)/ (*precision + recall*), on the task of identifying VOC vectors drawn from the diseased distribution in the test set. The omniscient model’s performance was retained as a comparison point for all subsequent models.

OBP multiplex sets were then evaluated on the same task and VOC population using the interface *f*(*P_X_*(*C*)) → *y*’, where *P_χ_* is the OBP multiplex sensing operation using candidate OBP set *X* as its input layer and *f* is the random forest model trained on the OBP multiplex’s outputs. For each proposed OBP set *X*, the F1 score of the combined model *f*(*P_X_*(*C*)) was produced.

### OBP Multiplex Optimization

To select a hero multiplex of OBPs, channels (i.e., OBP proteins) were selected one at a time to account for conditional relationships between OBPs. For instance, a nonanal-binding OBP may have high utility for detecting a disease but adding a second nonanal binder might be expected to provide minimal incremental benefit. Therefore, each OBP must be evaluated in the context of the other specific OBPs in an N-plex set. We built up the N-plex piecewise, so that sensor tests were conditioned on all preceding OBP choices. This operation was defined as follows:

Begin with the “hero sensor” and “best-yet sensor” comprising empty sets 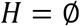 and 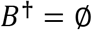 and *T* total experiments budgeted. Define the function *E*(*X*) to be the evaluation operation described above that returns an F1 score for a sensor comprised of OBPs *X*:

1. Randomly select *T/N* prospective sets of *N* – *len*(*H*) OBPs without replacement - this set of OBP sets is 5.
2. Evaluate each prospective set, retaining the highest performer: 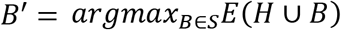
3. If *B*’ had a higher F1 score than *B*^†^, *B*^†^ ← *B*’
4. Select the OBP *o*’ that most reduces the sensor’s effectiveness when removed: 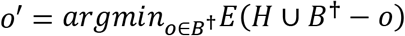
5. Append the new OBP to the hero sensor: 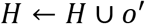 and *B*^†^ ← *B*^†^ – *o*’
6. Repeat steps 1-5 until *len*(*H*) = *N*

The intended output from this process was a set of *N* OBPs that maximize the F1 score on the classification task of identifying a disease by its VOC fingerprint. Optimization was performed on the Hyak supercomputer at the University of Washington, allowing about 100,000 potential multiplex sets to be evaluated per day on the disease classification task. In total, less than 1 million multiplex sets were evaluated to generate the designs presented here.

### Hyperparameter Optimization

All random forest and decision tree models and training behaviors used default hyperparameters except for the decision tree max depth, which was set to 1. No hyperparameter optimization was performed before or during model evaluation.

It was necessary to choose scaling values for *μ,* the VOC population average, and *v,* the disease scaling value, because present studies of COVID-19 VOCs are directional rather than quantitative - it was unknown to what degree VOC concentrations are perturbed in sick individuals. Qualitatively, perturbations between 2x and 10x appear likely^10^. This parameter choice was potentially high impact – a suitably small *v* renders the detection task impossible, while a large choice makes it trivial. Canine diagnostic studies provide performance baseline for what is possible in this space, with Chaber et al.^7^ observing diagnostic sensitivity in the 95% confidence interval of 93.1-97.6% and specificity within 95% confidence interval of 90.7%-100.0%. We extended dogs the generous assumption of omniscient VOC concentration detection and chose *μ* and *v* so that the omniscient baseline sensor would mimic the canine performance, achieving an F1 score in the range 0.93 to 0.97.

All experiments used M=5,000 simulated VOC populations, chosen to represent the upper end of realistic sample size in an early-phase human trial campaign. With this sample size, manual experimentation found that the naive choice of *μ = v =* 7, for a 2x increase in disease VOC levels, yielded an omniscient classifier 0.941 F1 score, which was in the target performance range above. The performance of the omniscient model then served as the baseline for evaluating sets of OBPs.

The falloff when introducing confounding disease signatures was steep, to F1~0.6 - the choice not to re-tune *v* allows for direct comparison between the two studies and emphasizes the challenge of this task with confounding diseases present. Because model performance is so sensitive to the choice of *v,* conclusions were drawn only from the relative performance of sensors to the omniscient baseline.

### Peptide Synthesis

Each chimeric peptide was synthesized using an automated solid-phase synthesizer (CS336X; CS-Bio, Menlo Park, CA, USA) through Fmoc-chemistry. In the reaction vessel, Wang resin (Novabiochem, West Chester, PA, United States), was treated with 20% piperidine in DMF to remove the preloaded Fmoc group. Incoming side chain protected amino acid was activated with HBTU (Sigma-Aldrich, St Louis, MO, USA) in dimethylformamide (DMF, Sigma-Aldrich), incubated with the resin for 45 min, then washed with DMF. This protocol was repeated for each subsequent amino acid. The synthesis reaction was monitored by UV absorbance at 301 nm. Following synthesis, the resulting resin-bound peptides were cleaved, and the sidechain deprotected using reagent- K (TFA:thioanisole:H2O:phenol:ethanedithiol (87.5:5:5:2.5), Sigma-Aldrich) and precipitated by cold ether. Crude peptides were purified by RP-HPLC with >95% purity (Gemini 10u C18 110A column). Peptide sequences were confirmed by MALDI-TOF mass spectrometry with reflectron (RETOF-MS) on an Autoflex II (Bruker Daltonics, Billerica, MA, United States).

### Probe Binding Validation

The two critical functions of the bi-functional chimeric peptides are both binding behaviors: they must bind selectively to a π-conjugated carbon surface to allow immobilization on a device and to their VOC targets. These are validated using SPR analysis and quartz crystal microbalance (QCM), respectively.

SPR (GE Healthcare, Biacore T200) was performed to validate specific adsorption of peptides onto a carbon substrate. Bare gold and carbon- and poly-L-lysine-functionalized substrates were used. Substrate fabrication used e-beam evaporation (CHA Industries, Solution Process Development System) of titanium (3-nm thick layer) then gold (47-nm thick layer) onto 1-cm^2^ optical quality glass. A 20-nm thick carbon layer was sputtered (Kurt J. Lesker Company, Lab 18 Sputter System) onto the fabricated gold substrates at a rate of 10 Å per minute, and poly-L-lysine coatings were drop-cast incubated with 0.1% v/v poly-L-lysine for 30 minutes and rinsed with diH2O, then air dried. A single-cycle kinetics curve was generated by sequential injection of MmedPep1-GBP peptide at 2-fold increasing concentrations up to 500 nM in buffer (100 mM KCl, 10 mM K+ phosphate). An injection flow rate of 30 μL/min and a temperature of 25 °C were controlled. Response values were plotted and fitted to a Langmuir isotherm curve by minimizing the sum of square differences between the dataset and the isotherm model.

VOC binding validation by QCM utilized chimeric peptides tethered to a substrate in a VOC- doped gas stream. Gold QCM substrates (Qsense, Phoenix, AZ, USA) were coated with 10 nm thick carbon film by evaporation (SPI, West Chester, PA, USA). 10 uL of 1 uM aqueous peptide was deposited and incubated for 1 hour at room temperature, and then the sample was dried in a nitrogen gas stream. A KSV model Z500 QCM (Qsense, Phoenix, AZ, USA) measured VOC binding to bare carbon (control), and peptide-coated substrates. QCM measurements were performed in an air stream bubbled through a DI water column, as shown in **Figure 4**. Humidified nitrogen was measured as the baseline, then VOC was introduced by bubbling nitrogen through the VOC/DI water mixture. The difference in resonant frequency between wet nitrogen and VOC mixture is reported - lower frequency indicates an increase in mass on the substrate due to gas molecules binding to the surface.

### Device Fabrication

CNT-FET devices were fabricated that enable peptide-sensitized detection of VOC targets from the vapor phase in a bottom gate configuration. Semiconducting single wall carbon nanotubes (SWCNT) (NanoIntegris) were chosen for their high on/off ratio and lower noise floor near and below the threshold voltage. The devices feature four spatially resolved sensing regions, each with eight transistors for 32 total sensors on each chip. Each region has a common source electrode and can be functionalized with a different VOC-binding chimeric peptide. CNT-FETs were incubated with chimeric peptide by pipette in a continuous flow humidified N_2_ environment. A photoresist-free, solution-deposition fabrication was used to minimize device hysteresis, which produced devices with transfer curve on-off ratios > 10^3^ and threshold voltages <5 V, as measured by a Keithley 2450 source meter. In brief, a back gate was made by reactive ion etching with CF4/CHF3 to remove the silicon dioxide layer on the backside of the silicon wafer, followed by deposition of chromium and gold. For the electrode contacts, photolithography then metallization and liftoff procedures were carried out. AZ1512 photoresist was then used to create channel banks in the sensing regions wherein CNT solution would be drop cast incubated prior to functionalization with chimeric peptide.

## Acknowledgments

Thank you to Kevin Jamieson for providing feedback on machine learning methodologies. This work was facilitated through the use of advanced computational, storage, and networking infrastructure provided by the Hyak supercomputer system at the University of Washington. Partial funding was received from WE-REACH program at the University of Washington through the RADx RAD program at NIDCR/NIH. Part of this work was conducted at the University of Washington Clean Energy Institute’s Clean Energy Testbeds, the Molecular Analysis Facility, and the Washington Nanofabrication Facility, a National Nanotechnology Coordinated Infrastructure (NNCI) site at the University of Washington, with partial support from the National Science Foundation via awards NNCI-1542101 and NNCI-2025489.

